# Neural Changes in Processing Visuo-Tactile Looming Stimuli Following Hand-to-Foot Sensorimotor Remapping

**DOI:** 10.64898/2026.08.25.746992

**Authors:** Matteo Girondini, Greta Madonna, Paolo Boffi, Alberto Gallace

## Abstract

Interactions with the environment follow stable spatial regularities that allow the brain to predict where sensory events are likely to occur. Although these expectations can adapt when actions repeatedly produce altered sensory consequences, whether spatial regularities learned through action influence subsequent sensory processing in the absence of action remains unclear. Participants underwent virtual reality (VR) sensorimotor remapping training in which right-hand interactions produced tactile feedback on either the ipsilateral (n=23) or contralateral foot (n=23). Feedback was either synchronous with hand–object contact, establishing a reliable action– sensation relationship, or asynchronous, providing comparable tactile exposure without a consistent temporal contingency. Before and after training, EEG was recorded during a visuo-tactile looming task in which participants passively observed objects approaching the hand while tactile stimulation was delivered to the hand (expected) or occasionally to the foot (unexpected). We examined the mismatch negativity (MMN) and P300 to determine whether the learned hand-to-foot regularity influenced subsequent processing of these events. P300 responses to hand stimulation increased selectively following synchronous training, indicating that learning a reliable hand-to-foot relationship altered subsequent processing of hand-related events outside the action context. In contrast, neither MMN nor P300 responses to foot stimulation differed between synchronous and asynchronous training, providing no evidence for direct transfer of the newly learned spatial mapping. Relative to baseline, foot-related responses instead showed contingency-independent changes consistent with more general exposure-related adaptation. Together, these findings show that spatial regularities learned through action can influence subsequent sensory processing beyond the context in which they are acquired, while highlighting constraints on their generalization across active and passive interactions.

**Significant Statements:** Our interactions with the environment rely on learned relationships between actions and their sensory consequences. But what happens when an action repeatedly produces sensation at an unusual location on the body? Using virtual reality, we trained participants to experience tactile feedback on the foot while interacting with objects using the hand and then measured brain responses during passive visuo-tactile stimulation. Learning a reliable hand-to-foot relationship altered subsequent neural processing of tactile events at the hand, even though the learned action was no longer performed. However, we found no evidence that the newly learned mapping directly transferred to the foot during passive observation. These findings show that sensorimotor learning can influence sensory processing beyond the context in which it is acquired, while revealing important constraints on how newly learned spatial relationships generalize across active and passive interactions

## Introduction

Our interactions with the environment follow stable spatial relationships between actions, external events, and their sensory consequences. Actions performed with different parts of the body typically produce sensory consequences at predictable locations: grasping an object with the hand generates tactile feedback on the fingers, while stepping on a surface produces feedback on the foot. Similarly, external events carry spatially predictable sensory consequences: a ball approaching the foot creates an expectation of tactile contact on the foot, rather than on another body part such as the hand. Through repeated experience, the brain learns these spatial regularities across both active and passive interactions with the environment and uses them to predict where sensory events are likely to occur (Jacquey et al., 2019). These learned spatial expectations, in turn, shape how sensory information is processed. During action, sensory consequences that match the predicted outcome are processed differently from unexpected consequences, as illustrated by the attenuation of self-generated touch (i.e., self-generated touch is typically perceived as less intense than externally generated touch; (Bays et al., 2006; Kilteni & Henrik Ehrsson, 2020)). Predictions can also arise independently of our own actions. Simply observing an external event approaching the body generates expectations about its likely sensory consequences: for instance, a visual stimulus approaching the hand facilitates the processing of tactile information at the expected location and modulates neural responses to anticipated contact (Cléry et al., 2015, 2017; Cléry & Ben Hamed, 2018; Kimura & Katayama, 2015, 2018, 2023). Whether generated through action or through observation of external events, learned spatial regularities allow the brain to anticipate where sensory information should occur and adjust its processing accordingly.

Notably, sensory expectations are not fixed but continuously adjusted as the regularities of our interactions with the environment change. When sensory information deviates from what is expected, the resulting prediction error signals a mismatch between current expectations and incoming sensory input (Friston, 2018; Pezzulo et al., 2022). When such deviations occur consistently, what initially constitutes a prediction error may therefore become a new regularity to be learned. This flexibility is well documented in sensorimotor interactions: repeated changes in the relationship between an action and its sensory consequences can recalibrate subsequent predictions (Krakauer & Mazzoni, 2011; Shadmehr et al., 2010). For example, delaying tactile feedback from an action initially reduces sensory attenuation, but repeated exposure to a fixed delay shifts the attenuation window toward the newly learned timing (Kilteni et al., 2019, 2023). Critically, such learning can include changes in where the sensory consequences of an action are expected to occur. Virtual reality provides a unique means to manipulate these spatial regularities, allowing the sensory consequences of an action to be systematically relocated to body locations where they would not naturally occur—for example, pairing a hand action with tactile feedback on the foot (Girondini, Bertoni, et al., 2025; Girondini, Montanaro, Lega, et al., 2024). However, it remains unclear how far these newly learned spatial expectations extend: do they remain specific to the action–sensation relationship in which they were acquired, or do they transfer to how sensory events are subsequently predicted in the absence of action?

To address this question, we tested whether learning a novel spatial relationship between an action and its tactile consequence influences subsequent neural processing in a passive context governed by an established spatial expectation. Participants underwent a VR sensorimotor remapping training in which right-hand interactions produced tactile feedback on either the ipsilateral or contralateral foot (between-group), creating a novel hand-to-foot action–sensation mapping. Critically, feedback was either synchronous with hand–object contact, establishing a reliable temporal relationship between action and sensory consequence, or asynchronous, providing comparable tactile exposure without a consistent action–feedback contingency (within-group). This comparison controls for distinct effects associated with learning a reliable sensorimotor relationship from nonspecific effects of repeated foot stimulation.

We then examined whether experience with this novel mapping influenced neural responses when sensory expectations were subsequently generated without action. At baseline and after each training condition, participants completed a visuo-tactile looming task while electroencephalography (EEG) was recorded. Participants passively observed virtual objects approaching their hand. Such looming stimuli provide spatial and temporal information about impending body contact and facilitate tactile processing at the predicted location and time (Cléry et al., 2015, 2017; Kimura & Katayama, 2015). Most approaching objects were followed by tactile stimulation of the hand, consistent with the established spatial relationship between an approaching object and its expected location of contact, whereas occasional tactile stimulation of the foot violated this relationship. Although this design shares features with a somatosensory oddball paradigm, with frequent hand standards and rare foot deviants (Shen et al., 2018a, 2018b), the sensory context was additionally defined by the visually specified location of impending contact.

We examined event-related potentials (ERPs) associated with expected and unexpected sensory events, focusing on the mismatch negativity (MMN), an early response associated with the detection of deviations from regular sensory patterns (Grundei et al., 2025), and the P300, which reflects later stages of stimulus evaluation and is sensitive to salience, attentional allocation, as well as multisensory integration for visuo-tactile stimuli (Polich, 2007; Kimura & Katayama, 2015). These components therefore provide complementary measures of early deviance and high-level-related sensory processing.

If spatial regularities acquired through action influence subsequent processing outside the sensorimotor context in which they are learned, hand-to-foot remapping should modulate neural responses to hand and/or foot stimulation during the passive looming task. A direct transfer of the newly learned spatial mapping would predict contingency-specific changes in responses to the remapped foot, whereas changes in hand-related responses could indicate that learning an alternative action–sensation relationship modifies the processing of the established hand-centered spatial expectation. Critically, comparing synchronous and asynchronous training allowed us to determine whether any such modulation depended on learning a reliable action–feedback relationship rather than on repeated tactile stimulation or task exposure.

## Materials and Methods

### Participants

We aimed to recruit 20 participants per group, consistent with previous EEG studies on sensorimotor training and tactile remapping (Girondini, Bertoni, et al., 2025; Soto-Faraco & Azañón, 2013). A total of 56 participants were initially enrolled and equally assigned to two groups: Ipsilateral Foot (IF) (n = 28, M age = 23.07 years, SD = 2.58, 21 females) and Contralateral Foot (CF) (n = 28, M age = 21.92 years, SD = 2.37, 18 females). Ten participants were excluded from EEG analyses due to poor data quality, resulting in a final sample of 46 participants (IF: n = 23; CF: n = 23). Participants were randomly assigned to groups. All provided written informed consent and completed a demographic questionnaire assessing age, gender, and handedness. The study was approved by the Ethics Committee of the University of Milano-Bicocca (Protocol No. 722) and conducted in accordance with the Declaration of Helsinki. Data and analysis code are publicly available at https://osf.io/bk5q3

### Experimental procedure

Each experimental session lasted approximately 1.5 hours. Participants were seated comfortably, fitted with a 64-channel EEG cap, and equipped with two vibrotactile actuators (3-v-coin shape actuators), one on the right hand and one on the assigned foot. They then wore a VR headset and were immersed in a virtual environment featuring an empty room with real-time hand tracking.

The experiment consisted of two paradigms: a sensorimotor remapping training, in which the spatial location and temporal contingency of somatosensory feedback during visuomotor interaction were manipulated, and a visuo-tactile looming task, administered before and after training to assess training-related changes in predictive processing. The experimental design included one between-subjects factor, feedback location (ipsilateral foot [IF] vs. contralateral foot [CF]), and one within-subject factor, temporal contingency (synchronous vs. asynchronous training) (Figure 1).

**Figure 1:**
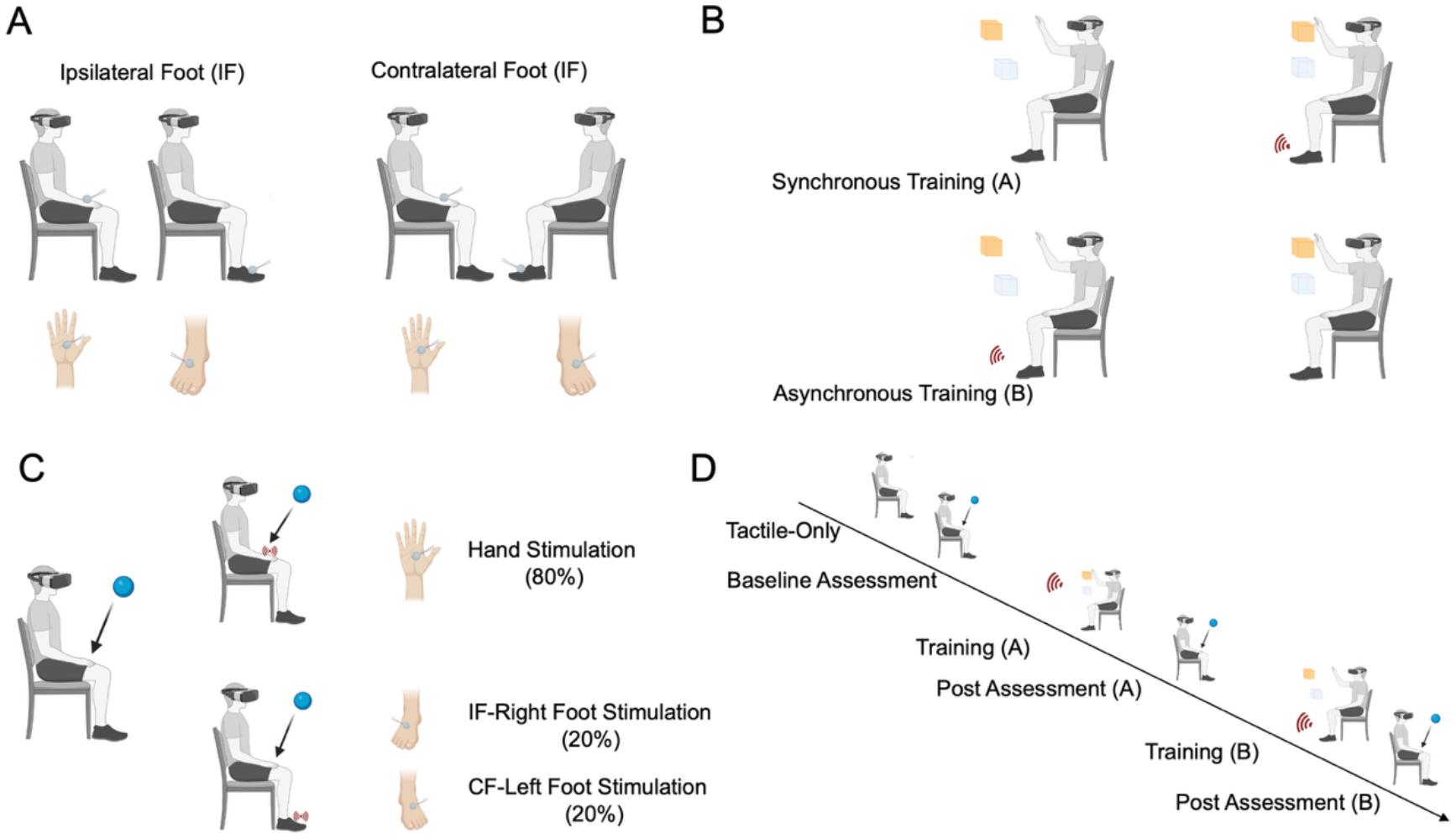
Experimental Design. A, Body locations used for tactile feedback in the Ipsilateral Foot (IF; right foot) and Contralateral Foot (CF; left foot) groups. B, Sensorimotor remapping training. Participants used the right hand to move a virtual cube toward a target. In the synchronous condition, vibrotactile feedback was delivered to the assigned foot at hand–cube contact; in the asynchronous condition, feedback occurred at random intervals independently of hand–cube contact. C, Visuo-tactile looming task. Virtual spheres approached the right hand and tactile stimulation was delivered at visual contact to the right hand on standard trials (80%) or to the assigned foot on deviant trials (20%). D, Experimental timeline. The pre-training baseline assessments were followed by synchronous and asynchronous training, each immediately followed by a visuo-tactile looming assessment. Training order was counterbalanced across participants.

The session began with two baseline assessments, both conducted before sensorimotor training. First, we recorded unimodal tactile evoked potentials while participants rested with their palms facing upward, and no visual stimuli were presented. Tactile stimulation was delivered separately to the hand and foot (100 trials per body part), with a 1-min rest between recordings. The order of hand and foot stimulation was counterbalanced across participants. Participants then completed a baseline assessment of the visuo-tactile looming task. They passively observed virtual spheres approaching their right hand while tactile stimulation was delivered either to the hand (standard trials, 80%) or to the foot (deviant trials, 20%). Each block consisted of 400 trials, with no more than one deviant occurring within five consecutive trials. No response to the tactile stimuli was required. To maintain attention, participants performed a cover task in which they mentally counted the number of blue spheres, irrespective of tactile stimulation location. Together, these pre-training assessments provided a reference for characterizing tactile responses in isolation and during visuo-tactile looming. Comparing the two allowed us to distinguish neural responses driven by somatosensory stimulation itself from those emerging when the same tactile input occurred within a visuo-tactile context that generated spatial expectations about impending contact. After completing the baseline assessments, participants rested for 5 min without the VR headset.

The session then continued with two experimental blocks, each comprising a sensorimotor remapping training phase followed immediately by a post-training visuo-tactile looming assessment The sensorimotor remapping paradigm was adapted from protocols previously used in our studies (Girondini, Bertoni, et al., 2025; Girondini, Montanaro, & Gallace, 2024; Girondini, Montanaro, Lega, et al., 2024; Girondini, Saccone, et al., 2025). During training, participants used their right hand to move a colored cube toward a semi-transparent target cube until contact was achieved. Upon contact, both cubes disappeared, and a new pair appeared. Participants completed two 10-min training conditions that differed in the temporal contingency between hand–cube contact and vibrotactile feedback. In the synchronous condition, vibrotactile stimulation was delivered to the foot at the moment of contact, establishing a consistent relationship between the action and its sensory consequence. In the asynchronous condition, vibrotactile stimulation was delivered at random intervals, independently of participants’ actions, disrupting this relationship (Figure 1B). Each training phase was immediately followed by the visuo-tactile looming task described above, allowing us to assess changes relative to the pre-training baseline. The order of the synchronous and asynchronous conditions was counterbalanced across participants, with a short break between experimental blocks. At the end of the session, participants were debriefed about the study aims.

### Hardware and Software

The VR setup consisted of a Meta Quest 2 head-mounted display (HMD) with a resolution of 1920 × 1832 pixels per eye. The HMD was connected via Oculus Link to an Asus ROG Strix laptop, featuring an AMD Ryzen 9 5900HX processor, 32 GB of RAM, an NVIDIA GeForce RTX 3080 graphics card, and a 17.3-inch display with a resolution of 1920 × 1080 pixels. The virtual environment was developed using the Unity 3D graphics engine In the visuo-tactile looming task and the synchronous condition of the training, the actuators were powered by an Arduino Uno board (5 V) connected to the VR laptop and temporally synchronized with virtual events using the Uduino library. For the asynchronous stimulation condition, a custom script was implemented to randomly activate the foot actuators. Activation intervals ranged from 0.5 to 4 seconds, with random delays of 1 to 4 seconds. Vibrotactile actuators were attached to the target body location (right hand or left/right foot) with adhesive tape.

### EEG Recording, Preprocessing, and Analysis

EEG data were recorded continuously using a 64-channel ActiCap system (Brain Products). Horizontal and vertical electrooculograms (EOGs) were recorded using electrodes placed below the left eye and at the outer canthi of the right eye. Electrode impedances were kept below 20 kΩ, and two external electrodes served as the ground reference. EEG and EOG signals were sampled at 5000 Hz and stored for offline analysis.

Preprocessing was performed using EEGLAB (Delorme & Makeig, 2004) and the FieldTrip toolbox (Oostenveld et al., 2011). Continuous data were band-pass filtered between 1 and 40 Hz and downsampled to 512 Hz. Channels exhibiting excessive noise were removed. Eye blinks and stimulation-related artifacts were corrected using independent component analysis (ICA). For each trial, stimulus-locked epochs were extracted from −100 ms to 500 ms relative to tactile stimulation onset and visually inspected to remove residual artifacts. Data were then re-referenced to the average of all remaining channels, and ERPs were baseline-corrected using the 100-ms pre-stimulus interval

Analyses focused on 32 fronto-central, central, centro-parietal, parietal, and parieto-occipital electrodes (F1, Fz, F2, F3, F4, F5, F6, FC1, FCz, FC2, FC3, FC4, C1, Cz, C2, C3, C4, CP1, CPz, CP2, CP3, CP4, P1, Pz, P2, P3, P4, P5, P6, PO3, POz, PO4), selected to capture the components of interest while avoiding channels affected by HMD-related noise

Because tactile stimulation was applied to different spatial locations across groups (right vs. left foot), analyses were first conducted separately for the IF and CF groups using paired-sample cluster-based permutation tests (Maris & Oostenveld, 2007). This approach allows for simultaneous statistical inference across both time points and electrodes, controlling for multiple comparisons while accounting for the spatiotemporal correlation structure of EEG data, thereby increasing sensitivity to detect genuine effects without inflating the false-positive rate. When a significant within-group effect was found, we tested whether its magnitude differed between groups by computing participant-level difference scores (i.e., synchronous minus asynchronous) and comparing these between the IF and CF groups.

The analysis focused on two ERP components associated with the processing of expected and unexpected tactile events: the mismatch negativity (MMN), elicited by deviant foot stimulation and typically observed 100–200 ms after stimulus onset, and the P300, elicited by both hand and foot stimulation and typically observed between 200 and 450 ms. To characterize these components in our visuo-tactile paradigm, we first compared responses to unimodal tactile stimulation with those recorded during the pre-training visuo-tactile looming assessment. This comparison allowed us to determine how tactile responses were modulated when stimulation occurred in the context of a visual stimulus approaching the participant’s hand.

The primary analysis tested the effect of sensorimotor training by comparing ERP responses following synchronous and asynchronous training within each group. This contrast was chosen for two reasons. First, the order of synchronous and asynchronous training was counterbalanced across participants, whereas the baseline assessment always occurred first and could therefore be affected by order or habituation. Second, both training conditions involved the same foot stimulation but differed in whether this stimulation was temporally contingent on the participant’s actions. Thus, the synchronous–asynchronous contrast isolates the contribution of action–feedback contingency, rather than the effect of repeated tactile stimulation alone.

As an additional control analysis, we compared ERP responses after each training condition with the pre-training visuo-tactile baseline. This analysis was used to assess whether changes could instead reflect nonspecific effects of repeated foot stimulation, task exposure, or habituation. Together, the primary and baseline comparisons allowed us to distinguish changes specifically associated with learning the action–feedback contingency from more general changes occurring over the course of the experiment. The same component-specific time windows and electrode set were used for the primary and baseline-related cluster-based permutation analyses. For visualization, ERP waveforms and mean amplitudes were displayed at representative electrodes within the corresponding significant clusters.

## Results

### Comparison of Training Performance in the IF and CF Groups

As a manipulation check, we assessed whether task performance differed between the synchronous and asynchronous training conditions or between the IF and CF groups. Performance was quantified as the number of trials completed during each 10-min training period. A 2 × 2 mixed ANOVA was conducted, with training condition (synchronous vs. asynchronous) as the within-subjects factor and group (IF vs. CF) as the between-subjects factor. The analysis revealed no significant main effect of training condition, F(1, 45) = 2.655, p = .110; no main effect of group, F(1, 45) = 0.026, p = .870; and no training condition × group interaction, F(1, 45) = 0.501, p = .647. Thus, participants completed a comparable number of trials across training conditions and groups, with an average of 118 trials (SEM = 1.5) per 10-min training period.

### ERP Responses to Visuo-Tactile Looming

To characterize the ERP components associated with the visuo-tactile looming task, we compared responses to tactile stimulation presented alone (unimodal condition) with responses to the same body locations during the baseline visuo-tactile looming task. This comparison allowed us to identify how tactile-evoked responses were modulated when stimulation occurred in the context of a visual stimulus approaching the hand.

For hand stimulation, a significant difference between unimodal and visuo-tactile conditions emerged in the P300 time window in both groups (IF: 200–450 ms, p < .001; CF: 200–450 ms, p < .001). The effect showed a widespread fronto-central and parietal distribution with a similar topography across groups. P300 amplitude was greater in the visuo-tactile than in the unimodal condition, indicating enhanced processing of hand stimulation when it coincided with an approaching visual stimulus (Figure 2, top).

**Figure 2:**
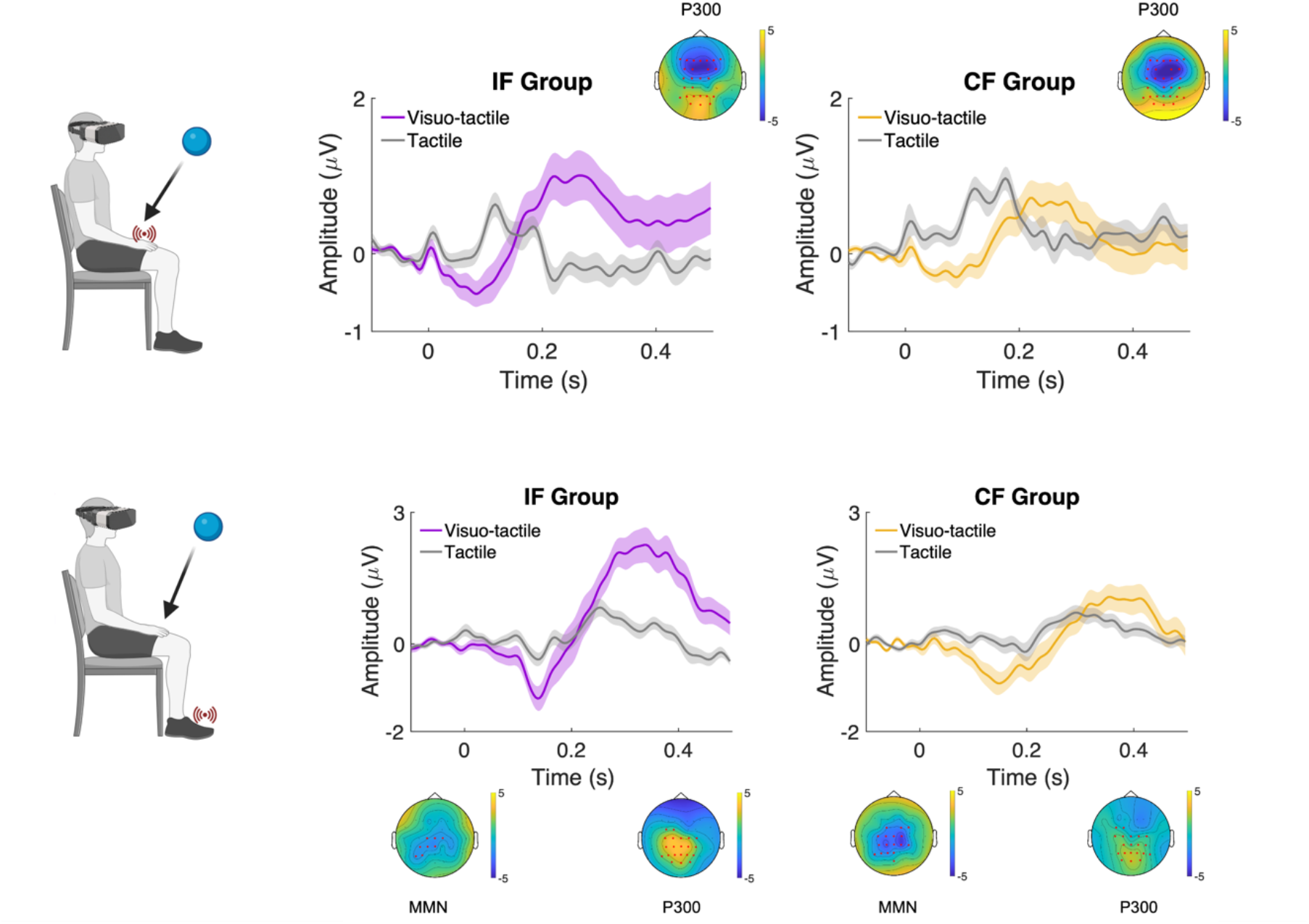
ERP responses to tactile stimulation during visuo-tactile looming. Grand-average ERP responses to hand (top) and foot (bottom) stimulation presented during unimodal tactile stimulation and the baseline visuo-tactile looming task, separately for the IF and CF groups. Hand stimulation during looming elicited an enhanced P300 (200–450 ms) relative to unimodal tactile stimulation in both groups. Deviant foot stimulation during looming elicited an early negative response consistent with the MMN (100–200 ms), followed by a later P300 (200–450 ms). ERP waveforms are shown at Pz for hand stimulation and CPz for foot stimulation as representative electrodes from the corresponding significant clusters. Topographic maps show the spatial distribution of the corresponding effects.

For foot stimulation, representing the deviant location in the visuo-tactile task, two components emerged. First, a negative deflection around 150 ms, consistent with the MMN, differed significantly between unimodal and visuo-tactile conditions in both groups (IF: 120–170 ms, p = .036; CF: 100– 200 ms, p < .001). This effect was centered over central electrodes and was more spatially extended in the CF group, although its magnitude did not significantly differ between groups. Second, both groups showed a later P300 response (IF: 200–450 ms, p < .001; CF: 200–430 ms, p < .010). In contrast to the MMN, the P300 differed significantly between groups (250–380 ms, p = .023), with a larger response in the IF than in the CF group over central electrodes (Figure 2, bottom).

Together, these results identified distinct early (MMN) and late (P300) responses to tactile events presented within the visuo-tactile looming context, supporting the use of these components and corresponding time windows in the subsequent analyses of training-related effects.

### Effects of Sensorimotor Remapping on Hand-Related Visuo-Tactile Responses

We first examined whether sensorimotor remapping altered responses to expected hand stimulation by comparing the synchronous and asynchronous training conditions within the IF and CF groups.

In both groups, P300 amplitude was significantly greater following synchronous than asynchronous training (IF: 350–380 ms, p = .030; CF: 260–440 ms, p = .005). This effect was distributed primarily over parieto-occipital electrodes. The magnitude of the synchronous–asynchronous effect did not significantly differ between the IF and CF groups (p > .100), indicating a comparable modulation across feedback locations.

To determine whether this effect was specifically associated with synchronous training rather than nonspecific changes over time, we additionally compared each post-training condition with the pre-training baseline. In both groups, P300 amplitude following synchronous training was significantly greater than at baseline (IF: 250–450 ms, p = .001; CF: 250–450 ms, p < .001). In contrast, P300 amplitude following asynchronous training did not differ significantly from baseline (IF: p > .100; CF: p > .260) (Figure 3).

**Figure 3:**
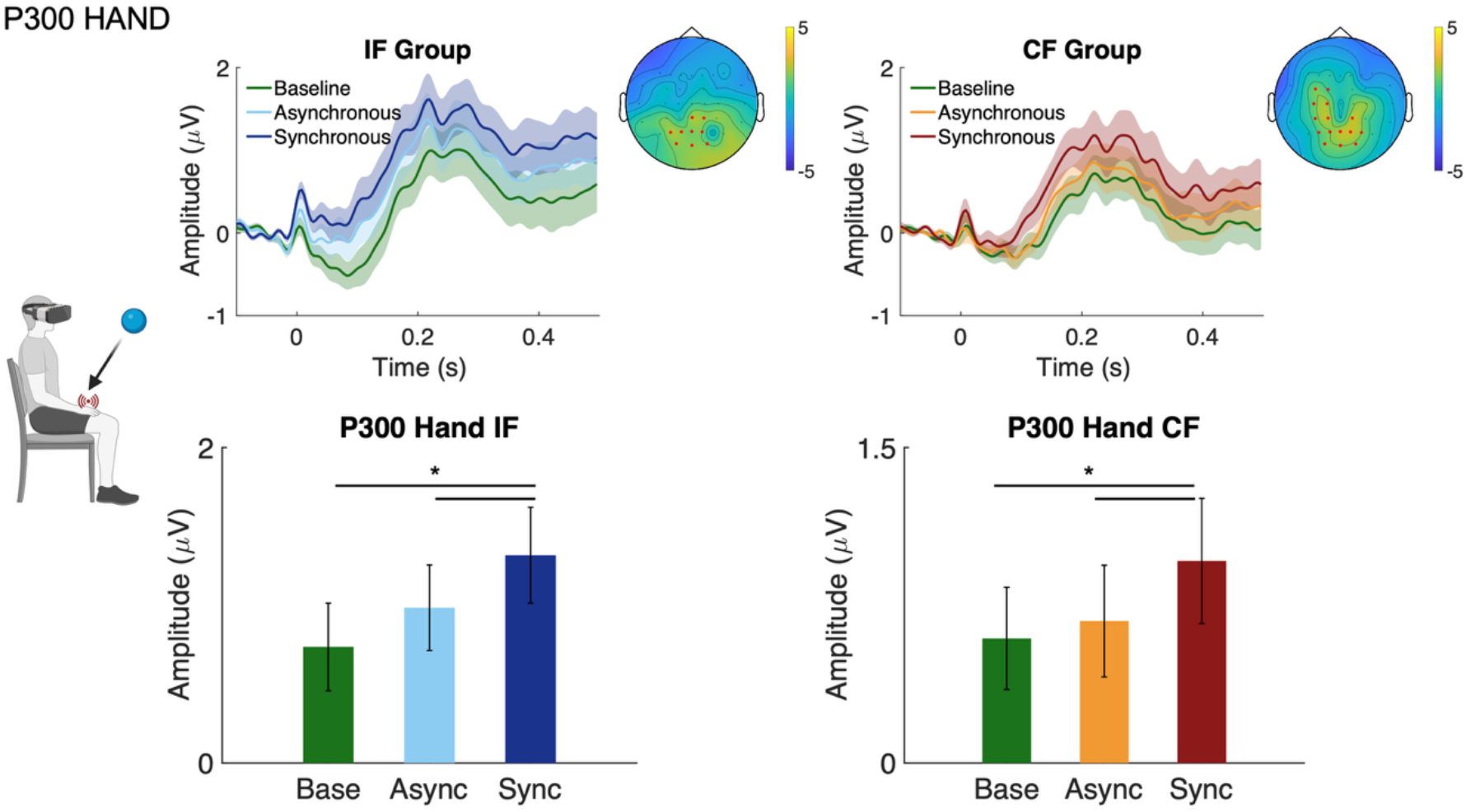
Contingency-dependent modulation of hand-related P300 responses following sensorimotor remapping. Grand-average ERP waveforms elicited by hand stimulation during the visuo-tactile looming task in the IF and CF groups at pre-training baseline and following asynchronous and synchronous training. Bar plots show mean P300 amplitudes across the three conditions. P300 amplitude was greater following synchronous than asynchronous training in both groups and was also greater following synchronous training than at baseline, whereas asynchronous training did not significantly differ from baseline. ERP waveforms and bar plots are shown at Pz as a representative electrode from the significant cluster. Shaded areas indicate SEM. Topographic maps show the spatial distribution of significant effects. p < .05.

Together, these results show that synchronous, but not asynchronous, sensorimotor remapping enhanced the P300 response to expected hand stimulation, with a comparable effect in the IF and CF groups. The absence of a change following asynchronous training suggests that this modulation cannot be explained by repeated tactile stimulation or task exposure alone, but instead depends on the temporal contingency between action and sensory feedback.

### Effects of Sensorimotor Remapping on Foot-Related Visuo-Tactile Responses

We next examined whether sensorimotor remapping altered responses to deviant foot stimulation, focusing on the MMN and P300 components.

For the MMN (100–200 ms), no significant differences were found between synchronous and asynchronous training in either group (IF: p > .999; CF: p = .158), providing no evidence that action– feedback contingency modulated the early response to deviant foot stimulation.

Comparisons with the pre-training baseline revealed a different pattern across groups. In the IF group, MMN amplitude was reduced following both synchronous (120–160 ms, p = .006) and asynchronous training (100–160 ms, p = .003), indicating that this reduction occurred independently of action–feedback contingency. In contrast, no significant changes from baseline were observed in the CF group following either synchronous (p = .151) or asynchronous training (p = .163) (Figure 4).

**Figure 4:**
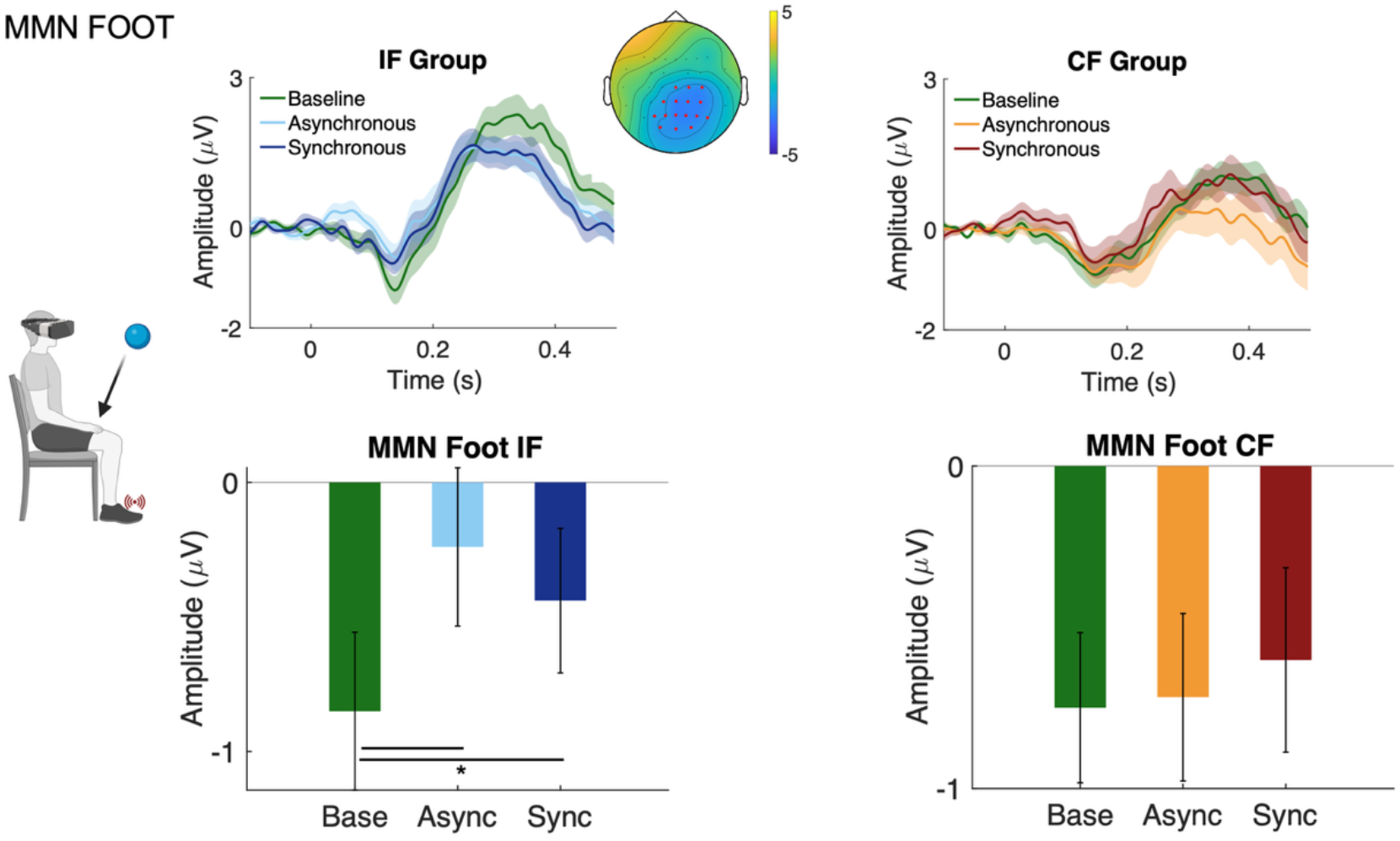
Contingency-independent changes in foot-related MMN responses following sensorimotor remapping. Grand-average ERP waveforms elicited by deviant foot stimulation during the visuo-tactile looming task in the IF and CF groups at pre-training baseline and following asynchronous and synchronous training. Bar plots show mean MMN amplitudes across the three conditions. No significant difference between synchronous and asynchronous training was observed in either group. In the IF group, MMN amplitude was reduced following both asynchronous and synchronous training relative to baseline, whereas no significant baseline-related changes were observed in the CF group. ERP waveforms and bar plots are shown at CPz as a representative electrode from the significant cluster. Shaded areas indicate SEM. Topographic maps show the spatial distribution of significant baseline-related effects. p < .05.

For the later P300 component (200–450 ms), no significant differences were observed between synchronous and asynchronous training in either group (IF: p = .388; CF: p > .076).

Comparisons with the pre-training baseline revealed reduced P300 amplitudes following both synchronous and asynchronous training in both groups (IF, synchronous: 280–450 ms, p < .001; asynchronous: 340–450 ms, p = .002; CF, synchronous: 340–450 ms, p = .002; asynchronous: 310–450 ms, p = .001) (Figure 5). Thus, the reduction in P300 amplitude occurred independently of action–feedback contingency.

**Figure 5:**
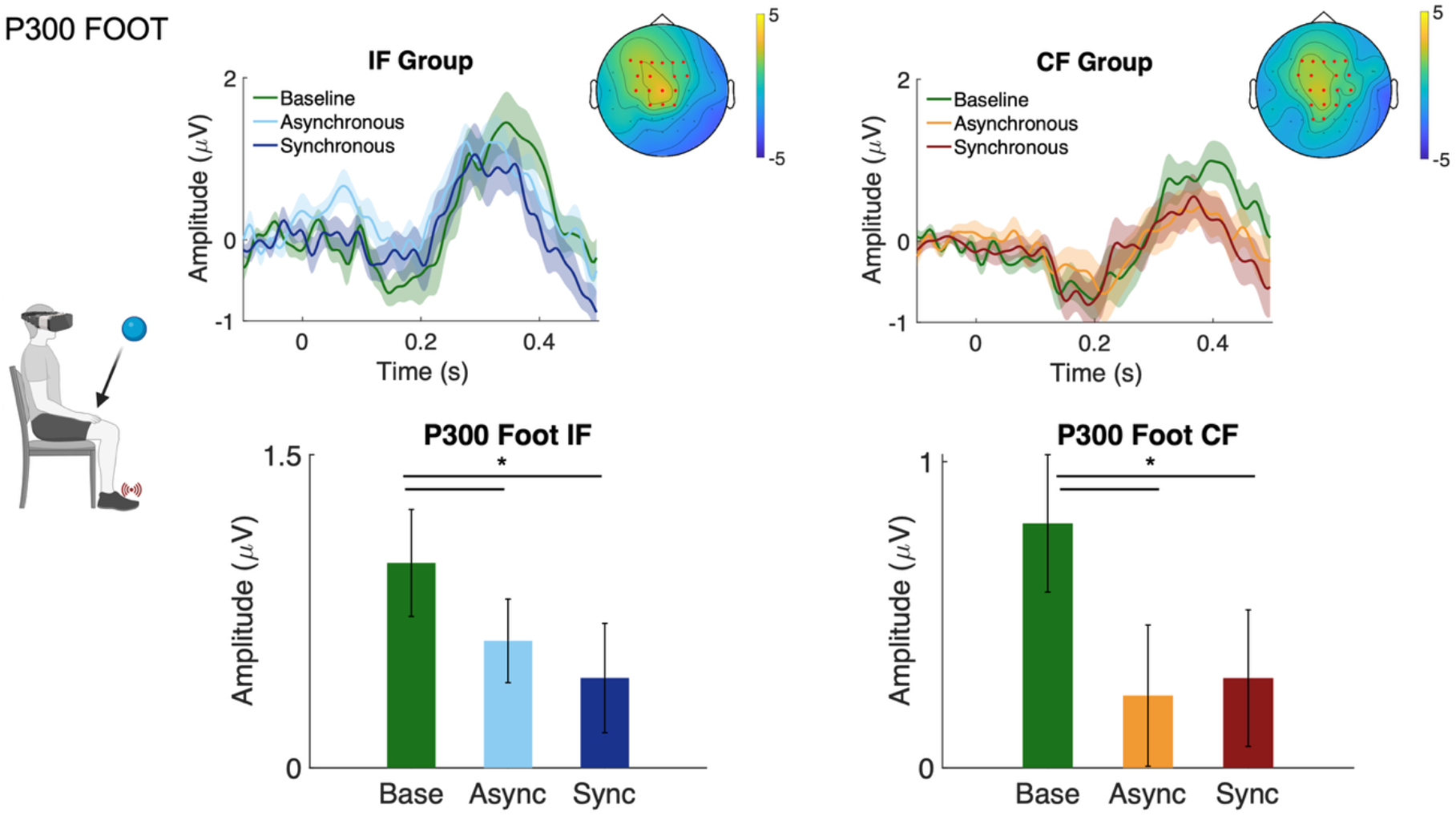
Contingency-independent reduction of foot-related P300 responses following sensorimotor remapping. Grand-average ERP waveforms elicited by deviant foot stimulation during the visuo-tactile looming task in the IF and CF groups at pre-training baseline and following asynchronous and synchronous training. Bar plots show mean P300 amplitudes across the three conditions. No significant difference between synchronous and asynchronous training was observed in either group. P300 amplitude was reduced following both asynchronous and synchronous training relative to baseline in both groups. ERP waveforms and bar plots are shown at Cz as a representative electrode from the significant cluster. Shaded areas indicate SEM. Topographic maps show the spatial distribution of significant baseline-related effects. p < .05.

Together, these results show that, unlike the P300 response to expected hand stimulation, responses to deviant foot stimulation were not differentially modulated by action–feedback contingency. Instead, both MMN and P300 amplitudes showed changes relative to baseline that were common to synchronous and asynchronous training. These contingency-independent effects differed across components: MMN attenuation was restricted to the IF group, whereas P300 attenuation was observed in both groups.

## Discussion

The present study investigated whether a novel spatial regularity learned through action can influence subsequent neural processing when that action is no longer performed. Participants underwent sensorimotor remapping in which right-hand interactions produced tactile feedback on either the ipsilateral or contralateral foot. Feedback was either synchronous with hand–object contact, establishing a reliable action–sensation relationship, or asynchronous, providing comparable tactile exposure without a consistent temporal contingency. We then assessed neural responses during passive observation of objects approaching the hand, with tactile stimulation delivered either to the naturally expected location (hand) or to the remapped location (foot). The results revealed a clear dissociation: P300 responses to hand stimulation increased selectively following synchronous training, whereas neither MMN nor P300 responses to foot stimulation differed between synchronous and asynchronous training. Thus, learning a reliable hand-to-foot relationship influenced subsequent neural processing outside the action context, but did not produce evidence for a direct transfer of the newly learned spatial mapping.

Visual looming stimuli approaching the body provide spatial and temporal information about impending contact, facilitating tactile processing at the expected location and time (Cléry et al., 2017; Kandula et al., 2017; Kimura & Katayama, 2015). Thus, in our looming task, virtual objects approaching the hand should primarily generate an expectation of tactile contact on the hand. Synchronous remapping training introduced a competing regularity: hand–object interactions were consistently followed by tactile feedback on the foot. If this newly learned mapping had transferred directly to the passive looming context, the approaching object might also have generated an expectation of foot stimulation, resulting in different neural responses to foot stimulation after synchronous compared with asynchronous training. However, neither the foot-related MMN nor P300 showed such a contingency-specific modulation. Instead, the effect of synchronous training emerged for stimulation at the naturally expected location, with an enhanced P300 to hand contact. This pattern suggests that learning the hand-to-foot mapping influenced how subsequent hand-contact events were processed, rather than replacing the pre-existing spatial expectation of contact on the hand with a new expectation of contact on the foot.

The enhanced P300 to hand-related sensory events may therefore reflect increased salience or higher-order processing when visual and tactile information converged again at the hand after learning an alternative spatial action–sensation regularity. This interpretation is supported by the broader role of the P300 as a saliency detector, also involved in attentional allocation and processing of visuo-tactile looming stimuli (Kimura & Katayama, 2015, 2023; Polich, 2007). For instance, stimuli that are inconsistent with spatial expectation elicit a larger P3 amplitude than congruent stimuli (Kimura & Katayama, 2015). In the present study, synchronous training may similarly have altered the significance of subsequent hand contact. Although the hand remained the naturally expected location of contact during looming, training had established the foot as a reliable sensory consequence of hand interaction. Consequently, tactile contact at the hand may have required greater evaluation in light of this recently acquired alternative regularity, resulting in an enhanced P300. Notably, this enhancement was comparable across the IF and CF groups, suggesting that it reflected the altered relationship between hand interactions and their sensory consequences rather than the specific spatial configuration of the feedback.

The MMN was nevertheless reduced relative to baseline following both training conditions in the IF group, but not in the CF group. Because this effect was independent of contingency, it may reflect adaptation to repeated foot stimulation rather than learning of the hand-to-foot relationship. Its restriction to the IF group raises the possibility that somatotopic organization contributed to early deviance processing, consistent with evidence that somatosensory MMN is sensitive to the cortical organization of stimulated body locations rather than simply their physical proximity (Lindenbaum et al., 2021; Shen et al., 2018b). This interpretation remains tentative, however, particularly in the absence of a significant between-group difference.

A similar contingency-independent pattern emerged for the later foot-related P300, which decreased relative to baseline following both training conditions and in both groups. This general reduction may reflect decreased novelty or salience following repeated exposure to foot stimulation, consistent with the sensitivity of P300 amplitude to stimulus repetition and habituation (Barry et al., 2020; Ravden & Polich, 1998). Importantly, repeated exposure alone cannot explain the hand-related P300 enhancement: hand stimulation was also repeatedly presented, yet its P300 increased selectively following synchronous training and remained comparable to baseline following asynchronous training. The opposing patterns therefore suggest a contingency-dependent modulation of hand-related processing against a background of more general exposure-related changes in responses to foot stimulation.

Overall, our findings suggest that spatial regularities learned through action can influence subsequent neural processing beyond the sensorimotor context in which they are acquired, but that this generalization is constrained. To our knowledge, this study represents one of the first attempts to test whether an experimentally induced cross-limb sensorimotor regularity can influence sensory processing outside the action context. Learning a reliable hand-to-foot relationship altered later processing of hand-related events without producing contingency-specific changes at the remapped location. This absence of direct spatial transfer should be interpreted cautiously, particularly because the assessment paradigm remained strongly centered on the hand: visual stimuli consistently approached the hand and predominantly predicted tactile contact at that location, potentially reinforcing the established hand-centered regularity and limiting expression of the newly acquired mapping. Moreover, because no behavioral measure of tactile perception was collected, the present findings demonstrate changes in neural processing rather than perceptual changes per se. Future studies should manipulate training duration, mapping stability, and the body location targeted during assessment, while combining neural and behavioral measures, to determine when newly learned spatial regularities generalize across active and passive contexts. Together, our findings show that newly learned sensorimotor regularities can influence subsequent sensory processing beyond the context in which they are acquired, while highlighting the constraints that govern their generalization across active and passive interactions.

## Acknowledgment

This paper was supported by a grant funded by the Italian Ministry of University and Research (MUR) – PRIN 2022, Grant n. 2022-NAZ-0172 to Alberto Gallace.

